# Structure of the *Saccharolobus solfataricus* type III-D CRISPR effector

**DOI:** 10.1101/2022.11.14.516469

**Authors:** Giuseppe Cannone, Dmytro Kompaniiets, Shirley Graham, Malcolm F White, Laura Spagnolo

**Author notes:** these authors contributed equally to the work (Giuseppe Cannone, Dmytro Kompaniiets). The Hormel Institute, University of Minnesota, Austin, MN 55912.

## Abstract

CRISPR-Cas is a prokaryotic adaptive immune system, classified into six different types, each characterised by a signature protein. Type III systems, classified based on the presence of a Cas10 subunit, are rather diverse multi-subunit assemblies with a range of enzymatic activities and downstream ancillary effectors. The broad array of current biotechnological CRISPR applications is mainly based on proteins classified as Type II, however recent developments established the feasibility and efficacy of multi-protein Type III CRISPR-Cas effector complexes as RNA-targeting tools in eukaryotes. The crenarchaeon *Saccharolobus solfataricus* has two type III system subtypes (III-B and III-D). Here, we report the cryo-EM structure of the Csm Type III-D complex from *S. solfataricus* (SsoCsm), which uses CRISPR RNA to bind target RNA molecules, activating the Cas10 subunit for antiviral defence. The structure reveals the complex organisation, subunit/subunit connectivity and protein/guide RNA interactions of the SsoCsm complex, one of the largest CRISPR effectors known.

## Introduction

CRISPR effectors are classified in two fundamental classes and six major types: Class 1 (multisubunit, types I, III and IV) and Class 2 (single subunit, types II, V and VI) (1,2). Type III CRISPR effectors are evolutionary related to type I systems and both share a CRISPR RNA (crRNA) binding “backbone” made up of Cas7 subunits. They differ in their large subunit: Cas8 for type I and Cas10 for type III (although Type I-D systems are hybrid, with a Cas10 subunit, and type III-E systems lack a Cas10 subunit (2)). Type III CRISPR effectors use their crRNA guide to bind cognate RNA molecules, which typically arise from mobile genetic elements (MGE) such as viruses that have previously been encountered by the CRISPR system. This binding activates the Cas10 subunit, which in different species harbours an HD nuclease domain for ssDNA cleavage, a cyclase domain for cyclic nucleotide synthesis, or a combination of both (reviewed in (3)). The cyclic nucleotides, which are composed of 3 to 6 AMP subunits linked by 3’, 5’ bonds (henceforth cA_3_, cA_4_ and cA_6_), act as second messengers to activate a wide range of accessory proteins. These include nucleases such as the Csx1/Csm6 family activated by cA_4_ or cA_6_ (4,5), the Can1/Can2/Card1 family activated by cA_4_ (6–8) and NucC activated by cA_3_ (9,10). These ancillary nucleases typically have relaxed specificity and target both viral and host nucleic acids to slow down infection or cause abortive infection (11). The coordination of anti-viral responses at the transcriptional level by cA_4_ are also possible (12).

In the past ten years, multiple structural studies have confirmed the overall “boot-shaped” structure of type III effectors, with Cas10 at the toe, Cas5 at the heel and helical constellations of Cas7 and Cas 11 subunits making up the shaft (or backbone) (13–21). The overall organisation resembles that of type I systems, and the two are sometimes known by the collective name “Cascade” (CRISPR-associated complex for antiviral defence) (22). Many groups have contributed to our current understanding of the target RNA recognition and activation of these effectors (reviewed in (3,23)), and type III effectors have been repurposed to develop sensitive new diagnostic assays (10,24,25). Despite this, the full diversity of type III systems has not been sampled at a structural level, and fundamental questions remain about the mechanism of activation of the Cas10 subunit on target RNA binding (21,26).

Here, we report the structure of the type III-D (SsoCsm) effector from the thermophilic crenarchaeon *Saccharolobus solfataricus*. SsoCsm was one of the first type III systems studied (13) and holds the record for the most unique subunits (eight) of any CRISPR effector. The structure of the SsoCSM complex bound to a 48 nt crRNA shows its architecture in unprecedented detail. This allows the appreciation of both stoichiometry and connectivity of the complex, which is unique in having a backbone composed of six Cas7-like subunits, encoded by four different genes.

## Materials and Methods

The SsoCsm complex was purified from *S. solfataricus* as described previously (13). Cryo-EM grids were prepared using an FEI Vitrobot Mark IV (Thermo Fisher) at 4°C and 95% humidity. A 4□μl volume of SsoCsm complex was applied to holey carbon grids (Quantifoil Cu R1.2/1.3, 300 mesh) covered by a graphene oxide layer (27), glow-discharged for 45□s at a current of 45□mA in an EMITECH K100X glow discharger. The grids were then blotted with filter paper once to remove any excess sample, and plunge-frozen in liquid ethane. All cryo-EM data presented here were collected on a ThermoFisher Titan Krios 300 microscope, equipped with a K2 direct detector, located at the eBIC facility. A total of 3,907 movies were collected in accurate hole centring mode using EPU. The MotionCorr and GCTF softwares from the Relion 3.1 image processing suite (28) were used for motion and CTF correction, respectively. Single particle analysis processing was carried out using the Relion 3.1 package, from corrected frames selection, manual particle picking to classification to generate templates for autopicking and subsequent 2D classification and 3D processing. The final reconstruction was obtained from 192,787 particles selected from classes representing both circular and elongated particles at a sampling rate of 1.046□Å per pixel and had an overall resolution of 3.52□Å, as calculated by Fourier shell correlation at 0.143 cutoff during post-processing. Alphafold2 models (29) built using ColabFold (30) were generated using the plugin implemented in the ChimeraX package (31). Individual subunits were fitted in the cryoEM map using the Dock in Map programme within the PHENIX 1.20.1 package (32). When more than one copy of a given subunit was present, the further subunits were fitted manually to produce a rigid body model of the fully assembled protein component of the complex.

## Results and discussion

### Cryo-EM structure of SsoCsm

We previously determined the structure of SsoCsm by negative staining TEM (13). The low resolution of negative staining techniques limited our ability to directly visualise the stoichiometry and connectivity of the subunits, as well as their interaction with the crRNA. We therefore used cryo-EM methods to elucidate the fine detail of this complex, gaining a deeper understanding of its molecular structure. Our previous negative staining and preliminary cryo-EM experiments allowed us to see that the particle distribution was lacking top and bottom views, however this didn’t impair the reconstruction of the assembly. On the other hand, cryo grids without a support would lead to preferential side views of the SsoCsm particles. We therefore decided to use graphene oxide (GO) coated grids to emulate the particle distribution typical of the carbon coated grids used in negative staining, at the same time as limiting the background that is typical even of very thin carbon films (33). Furthermore, in order to improve the particle contrast as well as maintaining resolution, we collected data in focus, inserting a Volta phase plate (34).

We solved the SsoCsm structure at a final overall resolution of 3.52 Å (Figure 1A and supplementary material). The comparison of the SisCmr complex shown in Figure 1A was edited to exclude the quite unusual Cmr7 subunit (35), which is present as 13 dimers decorating the crRNP particle (36). In contrast to the SsoCmr complex in negative staining (37), and other multi-subunit CRISPR systems such as he Type III-A *S. epidermidis* Csm complex (21), the SsoCsm complex did not undergo disassembly, showing that it is a rather stable assembly. The analysis of the local resolution of the map (Figure 1B), showing local resolutions in the range 3.411 to 6.101 Å, suggested that SsoCsm has some inherent flexibility, particularly with respect to the movement of the Cas10 catalytic subunit, which may be relevant for its activity (26). Overall, the structure has common features with other Csm and Cmr complexes, both bacterial and archaeal (Figure 1A). Poor resolution of the map at the position of the Cas10 subunit, visually shown in the resolution-filtered and colour-coded map shown in Figure 1B, made it quite difficult to model the entire complex therefore leaving doubts on the conformation of Cas10, in particular its N-terminal half.

**Figure 1.**
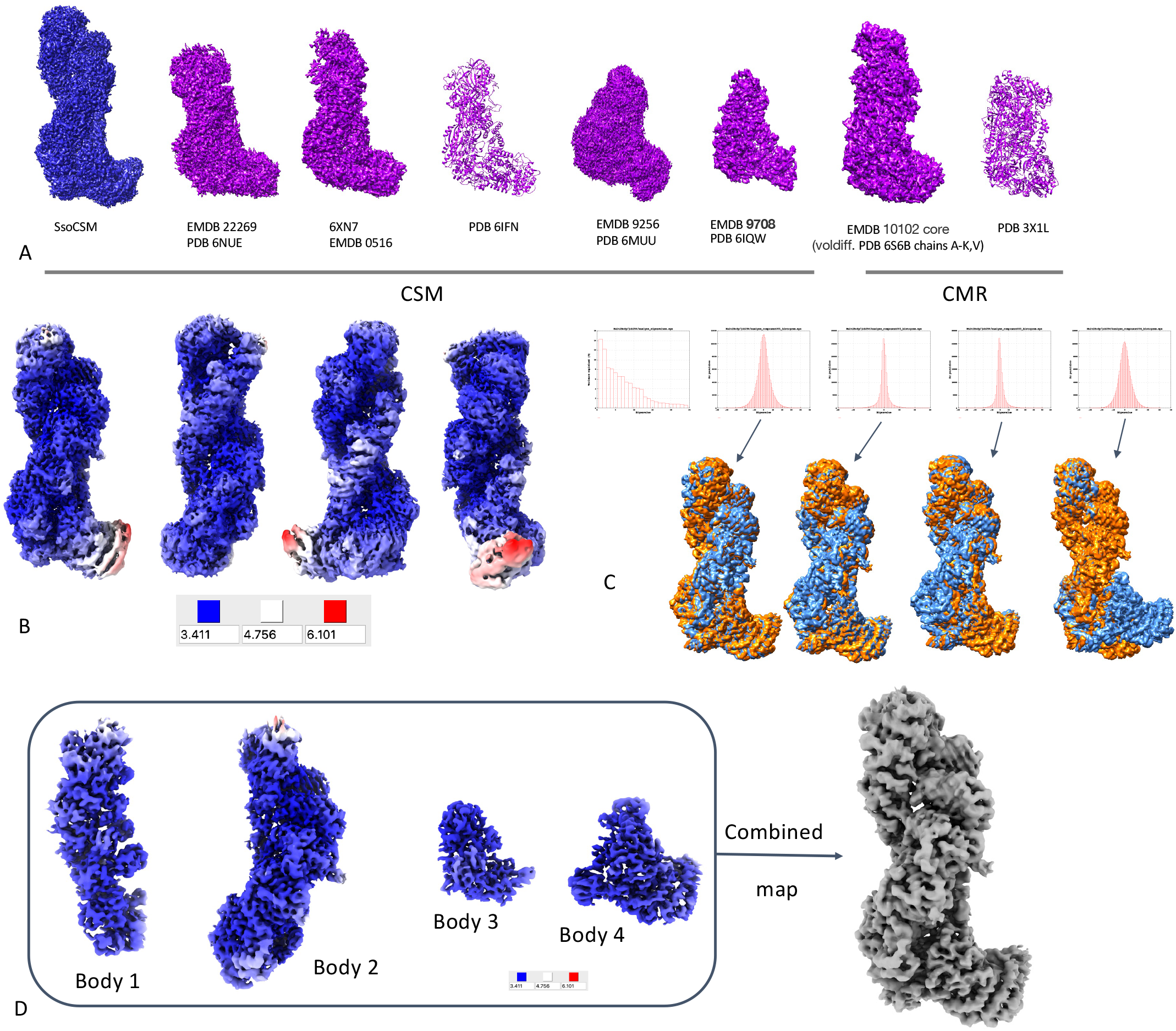
Structural analysis of SsoCSM. A. Comparison of the SsoCSM cryo-EM map with other published CSM and CMR complexes. B. Resolution-filtered SsoCSM map, colour-coded based on local resolution. C. Results of MultiBody analysis, showing histograms for the statistical analysis of eigenvectors 1-4 and volumes for these. The volumes for the two extremes of the analysis are coloured blue and orange, respectively, for eigenvectors 1 to 4. D. 3D reconstructions of bodies 1-4 in the MultiBody analysis, surface coloured based on the local resolution, and full map obtained combining the four bodies in Chimera.

A Multibody refinement experiment allowed the visualization of the main movements within SsoCsm, as shown in Figure 1C. We decided to group each strand of the double filament in one group (body 1 and body2), while the heel was analysed as body 3 and the tip as body 4. On analysing the output of this four-body refinement, eigenvalues 1 and 2 explain ~14 % and ~12 % of the variance respectively, while eigenvalues 3 and 4 account for around 8 % each (Figure 1C, histograms at the top). The maps at the extremes of the movements are shown in orange and blue in Figure 1C, lower panel, and movies 1-4 are in the supplementary material. The output from the first eigenvalue analysis highlights an overall swivel movement of body 1 on the long axis of the structure, suggesting an opening to expose the RNA backbone that may be relevant to accommodate target RNA binding. The output from the second analysis highlights an opening of body 2 on the x axis, suggesting an opening towards the crRNA 5’-handle. The third movement is again an opening of body 2, this time on the z axis, again allowing more accessibility towards the 5’ of the RNA molecule bound to the complex. The more complex movement in the fourth analysis is a large rotation+translation of the catalytic subunit (body 4), both on the x axis. The latter is likely the component that most affected the anisotropy of the overall reconstruction resolution.

The outputs from the Multi Body experiment in Relion 3.1 (28) were used to assemble a combined map using the Chimera software (31). As shown in Figure 1D, the local resolution of the individual bodies was much more homogeneous, leading to a resolution that would allow confident modelling even for the otherwise blurred catalytic subunit (Figure 2B and C; Figure 3B-D). The more detailed combined map assembled in Chimera was used for further fitting experiments, instead of the consensus map.

**Figure 2.**
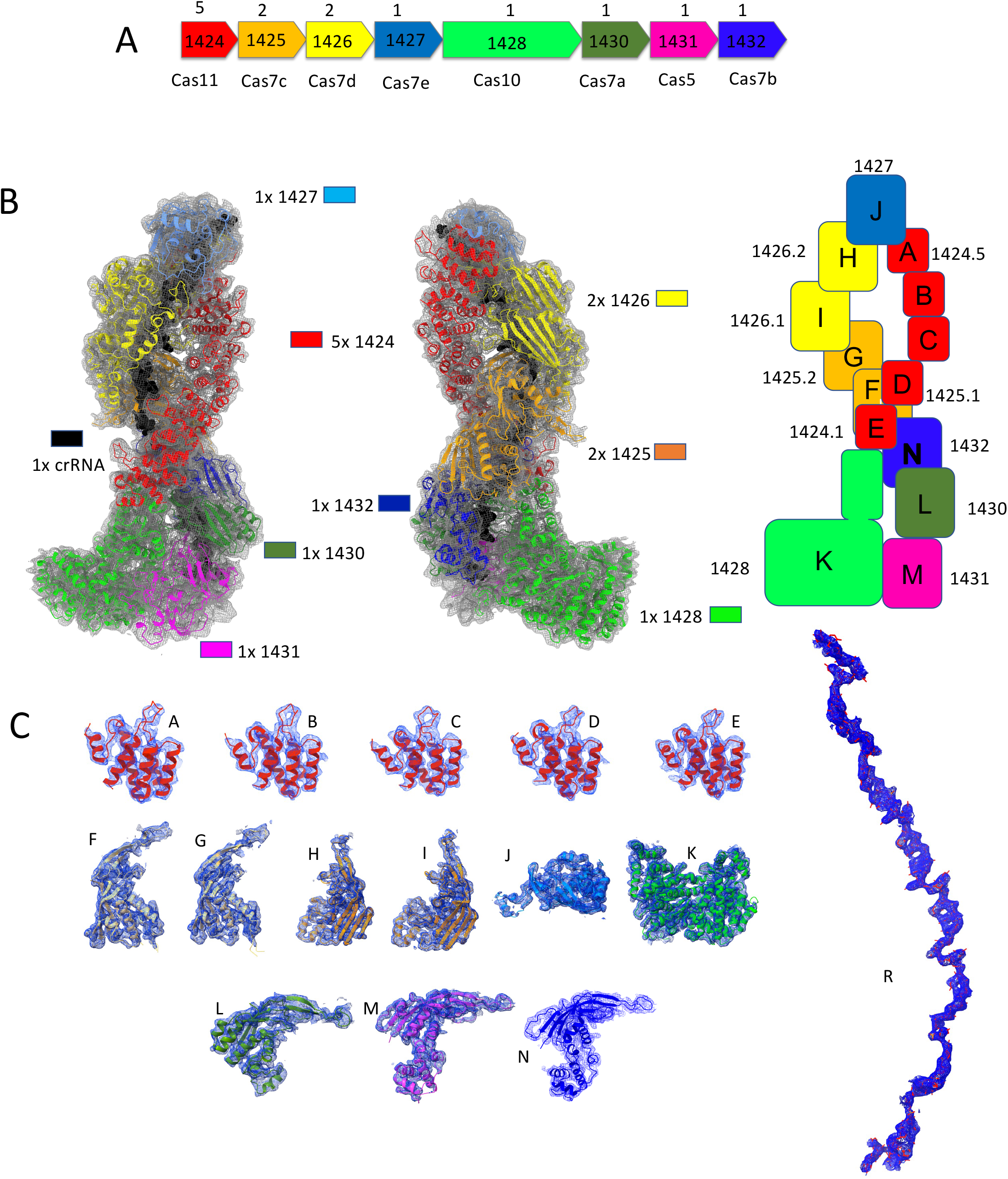
Interpretation of the SsoCSM cryo-EM structure. A. SsoCSM operon structure. ORF numbers are shown along their Cas identity and stoichiometry within the complex B. Fitting individual subunit in the cryo-EM map: front and back view are shown, as well as a schematic diagram summarizing position within the complex, stoichiometry and chain name. The RNA molecule (chain R) is shown as black spheres to emphasize its position within the ribonucleoprotein. C. Evaluation of the fit of each chain into the corresponding part of the cryo-EM map.

**Figure 3.**
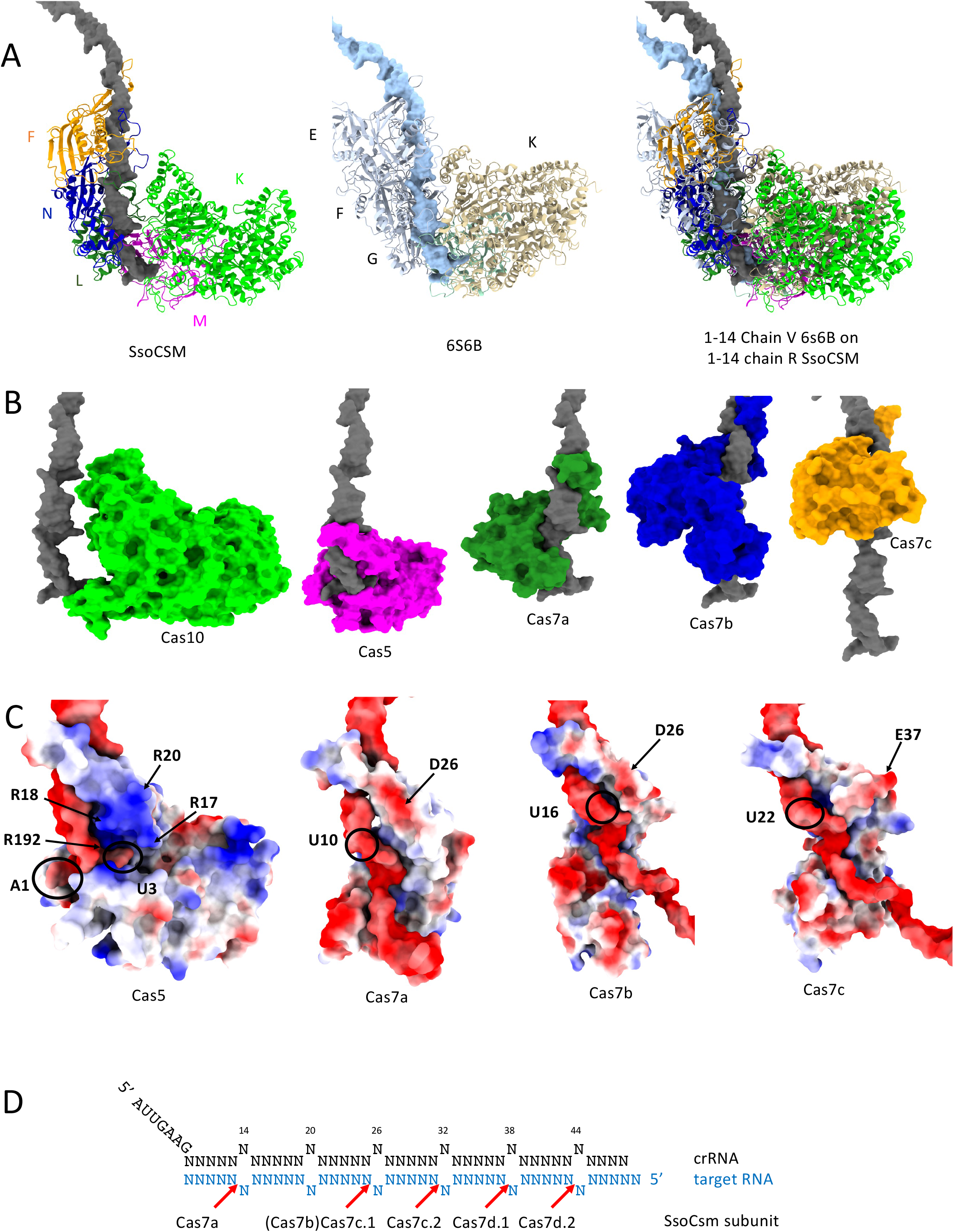
RNA threading within the SsoCSM cryo-EM structure. A. Four orthogonal views of SsoCSM, with the crRNA density shown red to highlight the position of the solvent-exposed RNA. B. Comparison of the 5’ region of crRNA is SsoCSM and SisCMR (pdb 6S6B), showing the distinct folding of the 5’ handle in Sso crRNAas. C. Chain K and L make close contacts with the 5’ crRNA handle.

### Fitting ColabFold models, model refinement and analysis

The operon structure of the SsoCsm system, comprising one cas11 gene (1424), one cas5 gene (1431), one cas10 gene (1428), and five cas7-like genes (1425, 1426, 1427, 1430 and 1432), is shown in figure 2A. The structures of each SsoCsm subunit were predicted using ColabFold (30) as built in the ChimeraX software package (31) (Supplementary Figure 1). Each model structure had a slightly different coverage and Local Distance Difference Test (lDDT) values (30,38), with 1432 showing the biggest discrepancies across models.

We fitted one copy of each model in the composite cryoEM map using the *Dock Predicted Model* programme from the PHENIX 1.20.1 package (32). Subunits present in multiple copies in the complex were assessed by visual inspection of the map and fitted manually. The rigid-body fitted model contained 5 copies of 1424 (chains A-E), 2 copies of 1425 (chain F,G), 2 copies of 1426 (chains H,I), 1 copy of 1427 (chain J), 1 copy of 1428 (chain K), 1 copy of 1430 (chain L), 1 copy of 1431 (chain M) and 1 copy of 1432 (chain N), as well as one 48-ribonucleotide-long RNA chain. These coordinates were refined using the PHENIX Real Space Refinement (32), Refmac-Servalcat (39) and Coot (40) packages, to obtain the model shown in Figure 2B. Figure 3C shows how each chain within the assembly fits in the corresponding part of the map.

This cryo-EM structural analysis yielded a model with an overall stoichiometry of Cas10_1_:Cas5_1_:Cas11_5_:Cas7_7_:crRNA_1_ and a calculated molecular weight of 428 kDa, in close agreement with the value of 423 kDa estimated by native mass spectrometry (13). The crRNA is 48 nt long, including the 8 nt 5’-handle. As the complex was purified from the native host, the bound crRNA is heterogeneous, being sampled from the over 200 spacers present in *S. solfataricus* P1 (13,41), so the 40 nt of the spacer have been modelled as uracils. Overall, the structure is reminiscent of that of other Csm and Cmr complexes (some examples are shown in Figure 1A), formed of two intertwined helical filaments of multiple stacked subunits wrapping around the crRNA, although SsoCsm is the tallest. Reconstitution experiments for SsoCsm (42) showed that each subunit was essential for RNase activity except for the 1427 subunit, which caps the structure (Figure 2B). Thus, the four different Cas7 subunit types making up the backbone are all essential and cannot substitute for one another.

Comparison with the “core” of SisCmr (pdb 6S6B, chains A-K, V) shows that the overall RNA conformation is quite similar, in particular at the 5’-handle (Figure 3A), and that despite some obvious structural differences in individual subunits the architecture supporting the RNA threading is equivalent in both complexes. The crRNA within SsoCsm has a standard conformation, bound to the intertwined Cas7 filament (Figure 3 and Figure 4B). Examination of the foot of the complex shows that there is almost no protein:RNA interaction for the Cas10 subunit, while Cas5 makes tight contacts with the guide RNA molecule (Figure 3B). These interactions are represented at finer detail in Figure 3C, which shows that the 5’ handle of the crRNA sits close to a patch of basic residues (R17, 18, 20 and 192) that are clustered together, spatially close to U3. The final nucleotide of the 5’ handle, G8 (sometimes also known as position −1 to discriminate between the handle (−8 to −1) and spacer (1 to X) regions of the crRNA), is flipped, as seen in other type III complexes such as SisCmr (36). Biochemical studies have demonstrated that base pairing with target RNA at this position prevents activation of Cas10 (43–45).

The remainder of the crRNA adopts a regular pattern with every sixth nucleotide adopting a flipped orientation (Figure 3). These positions correspond to the sites of cleavage of bound target RNA (46), which is catalysed by the Cas7-like backbone subunits (47–49).This activity is important for the dissociation of target RNA and deactivation of the Cas10 subunit (44,45). Previous studies showed that SsoCsm is unusual in not cleaving at one of the flipped sites, generating a spacing of 12, 6, 6, 6 between sites (42). The missed cleavage corresponds to the Cas7b subunit (1432; chain N in the PDB file, Figure 3C, D), while subunits competent for RNA cleavage are Cas7a (1430, chain L in the coordinates file), Cas7c (1425, chains F and G) and Cas7d (1426, chains H and I). It is not obvious from this apo structure why Cas7b does not cleave bound target RNA, as it has a similar structure to the other Cas7 subunits and a plausible active site aspartate residue, but this missed cleavage may be due to differences in local target RNA structure.

The diversity of Cas7 subunits in SsoCsm is a unique feature of the complex, as most type III systems make do with just one. It also seems to present some unique challenges for assembly of the complex, which must build in a strict order (from Cas5) with one Cas7a, one Cas7b, two Cas7c and two Cas7d subunits before capping the structure with Cas7e (Figure 2). The *in vitro* reconstitution experiments reinforce the requirement for each of these subunits in the active complex (42). The two duplicated subunits (7c and 7d) thus make different subunit contacts along the length of the backbone. This might be more easily achieved if these subunits are already dimeric in nature, but this has yet to be confirmed.

## Concluding remarks

Here, we have presented the cryo-EM Structure of the type III-D CRISPR effector SsoCsm. With eight different subunits and a molecular weight of 430 kDa, this is one of the largest and most complex CRISPR effectors studied to date. A unique feature is the complexity of the Cas7 backbone structure, which is assembled from seven subunits encoded by five genes. Analysis of the structure reveals conformational flexibility that most likely relate to target RNA capture and Cas10 activation. Unfortunately, the inherent diversity of crRNA in this complex precluded analysis of target bound states, but this is a promising area for future study.

## Acknowledgements

We acknowledge support from Dr Daniel Clare at the UK’s national Electron Bio-imaging Centre (eBIC) for the setup of the data collection. We acknowledge Margaret Mullin at the Gilmorehill electron microscopy facility Glasgow for support with initial sample screening. We acknowledge the Scottish Centre for Macromolecular Imaging (SCMI) for access to cryo-EM instrumentation for sample screening, funded by the MRC (MC_PC_17135) and SFC (H17007).

## Figure Legends

**Movies 1-4.**
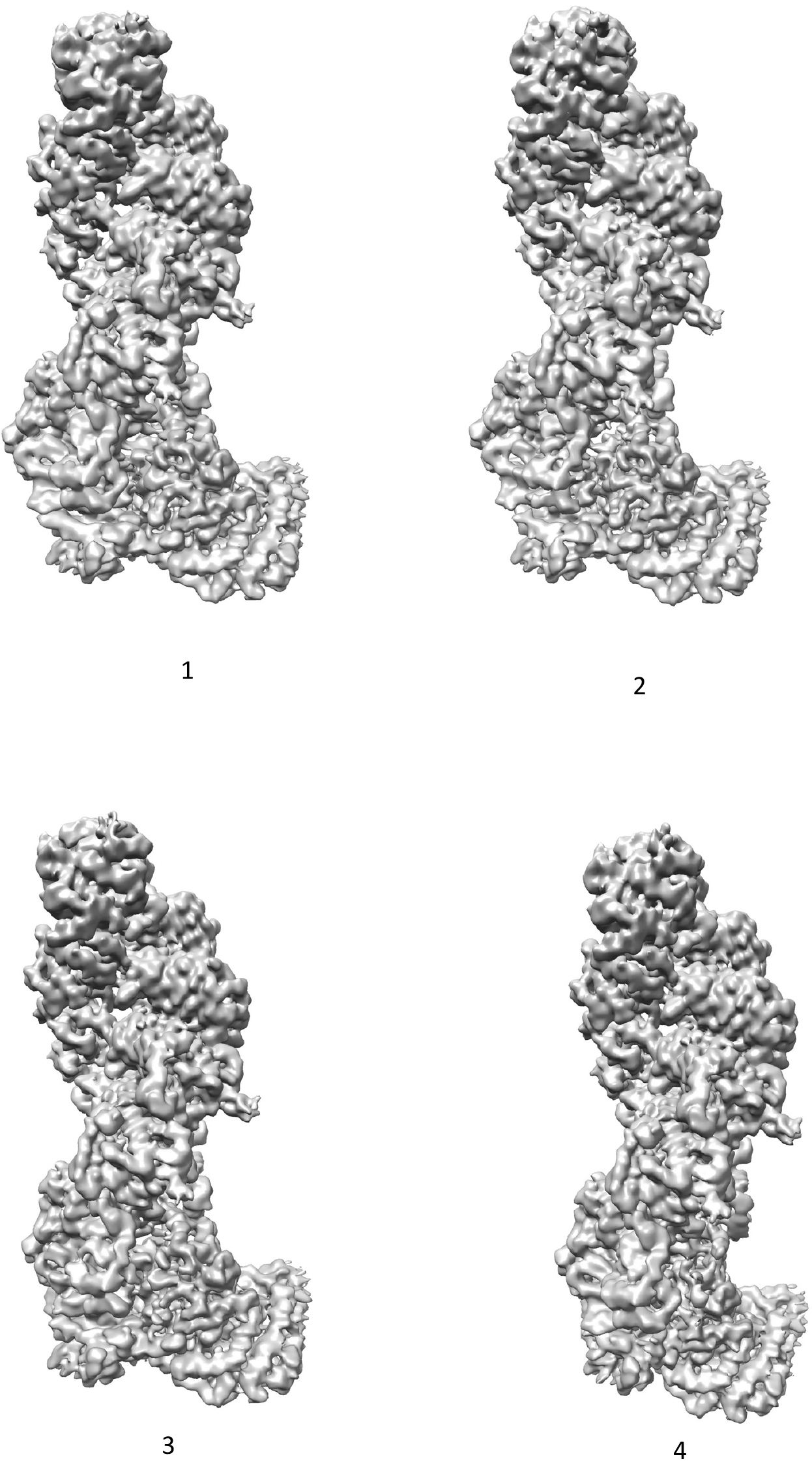
Movies for eigenvector analyses 1-4.

**Figure S1.**
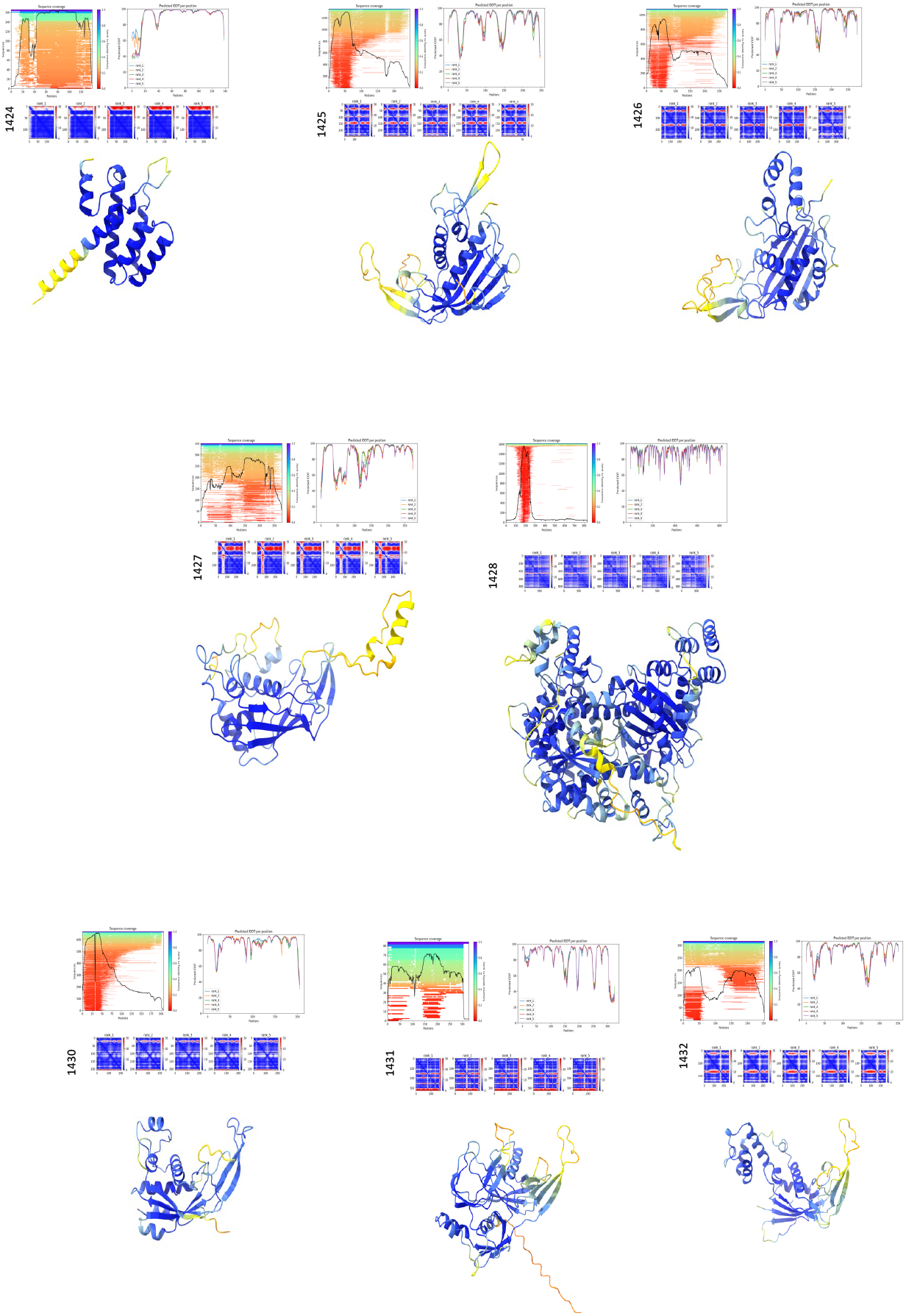
Structural predictions. AlphaFold2 models for the individual subunits. The data include lDDT diagrams, which are useful in assessing the local quality of the structure.

Table 1. **Cryo-EM data collection, refinement and validation statistics**.

## References

1. Makarova, K.S., Wolf, Y.I., Alkhnbashi, O.S., Costa, F., Shah, S.A., Saunders, S.J., Barrangou, R., Brouns, S.J., Charpentier, E., Haft, D.H. et al. (2015) An updated evolutionary classification of CRISPR-Cas systems. Nature reviews. Microbiology, 13, 722–736.

2. Makarova, K.S., Wolf, Y.I., Iranzo, J., Shmakov, S.A., Alkhnbashi, O.S., Brouns, S.J.J., Charpentier, E., Cheng, D., Haft, D.H., Horvath, P. et al. (2020) Evolutionary classification of CRISPR-Cas systems: a burst of class 2 and derived variants. Nature Rev Microbiol, 18, 67–83.

3. Tamulaitis, G., Venclovas, C. and Siksnys, V. (2017) Type III CRISPR-Cas Immunity: Major Differences Brushed Aside. Trends Microbiol., 25, 49–61.

4. Kazlauskiene, M., Kostiuk, G., Venclovas, C., Tamulaitis, G. and Siksnys, V. (2017) A cyclic oligonucleotide signaling pathway in type III CRISPR-Cas systems. Science, 357, 605–609.

5. Niewoehner, O., Garcia-Doval, C., Rostol, J.T., Berk, C., Schwede, F., Bigler, L., Hall, J., Marraffini, L.A. and Jinek, M. (2017) Type III CRISPR-Cas systems produce cyclic oligoadenylate second messengers. Nature, 548, 543–548.

6. McMahon, S.A., Zhu, W., Graham, S., Rambo, R., White, M.F. and Gloster, T.M. (2020) Structure and mechanism of a Type III CRISPR defence DNA nuclease activated by cyclic oligoadenylate. Nature communications, 11, 500.

7. Zhu, W., McQuarrie, S., Gruschow, S., McMahon, S.A., Graham, S., Gloster, T.M. and White, M.F. (2021) The CRISPR ancillary effector Can2 is a dual-specificity nuclease potentiating type III CRISPR defence. Nucl. Acids Res., 49, 2777–2789.

8. Rostol, J.T., Xie, W., Kuryavyi, V., Maguin, P., Kao, K., Froom, R., Patel, D.J. and Marraffini, L.A. (2021) The Card1 nuclease provides defence during type-III CRISPR immunity. Nature, 590, 614–629.

9. Lau, R.K., Ye, Q., Birkholz, E.A., Berg, K.R., Patel, L., Mathews, I.T., Watrous, J.D., Ego, K., Whiteley, A.T., Lowey, B. et al. (2020) Structure and Mechanism of a Cyclic Trinucleotide-Activated Bacterial Endonuclease Mediating Bacteriophage Immunity. Mol. Cell, 77, 723–733.

10. Grüschow, S., Adamson, C.S. and White, M.F. (2021) Specificity and sensitivity of an RNA targeting type III CRISPR complex coupled with a NucC endonuclease effector. Nucl. Acids Res.,gkab119.

11. Rostol, J.T. and Marraffini, L.A. (2019) Non-specific degradation of transcripts promotes plasmid clearance during type III-A CRISPR-Cas immunity. Nat Microbiol, 4, 656–662.

12. Charbonneau, A.A., Eckert, D.M., Gauvin, C.C., Lintner, N.G. and Lawrence, C.M. (2021) Cyclic Tetra-Adenylate (cA4) Recognition by Csa3; Implications for an Integrated Class 1 CRISPR-Cas Immune Response in Saccharolobus solfataricus. Biomolecules, 11.

13. Rouillon, C., Zhou, M., Zhang, J., Politis, A., Beilsten-Edmands, V., Cannone, G., Graham, S., Robinson, C.V., Spagnolo, L. and White, M.F. (2013) Structure of the CRISPR interference complex CSM reveals key similarities with cascade. Mol. Cell, 52, 124–134.

14. Spilman, M., Cocozaki, A., Hale, C., Shao, Y., Ramia, N., Terns, R., Terns, M., Li, H. and Stagg, S. (2013) Structure of an RNA Silencing Complex of the CRISPR-Cas Immune System. Mol. Cell, 52, 146–152.

15. Staals, R.H., Agari, Y., Maki-Yonekura, S., Zhu, Y., Taylor, D.W., van Duijn, E., Barendregt, A., Vlot, M., Koehorst, J.J., Sakamoto, K. et al. (2013) Structure and activity of the RNA-targeting Type III-B CRISPR-Cas complex of Thermus thermophilus. Mol. Cell, 52, 135–145.

16. Benda, C., Ebert, J., Scheltema, R.A., Schiller, H.B., Baumgartner, M., Bonneau, F., Mann, M. and Conti, E. (2014) Structural model of a CRISPR RNA-silencing complex reveals the RNA-target cleavage activity in Cmr4. Mol. Cell, 56, 43–54.

17. Jia, N., Mo, C.Y., Wang, C., Eng, E.T., Marraffini, L.A. and Patel, D.J. (2019) Type III-A CRISPR-Cas Csm Complexes: Assembly, Periodic RNA Cleavage, DNase Activity Regulation, and Autoimmunity. Mol. Cell, 73, 264–277 e265.

18. Liu, T.Y., Liu, J.J., Aditham, A.J., Nogales, E. and Doudna, J.A. (2019) Target preference of Type III-A CRISPR-Cas complexes at the transcription bubble. Nature communications, 10, 3001.

19. You, L., Ma, J., Wang, J., Artamonova, D., Wang, M., Liu, L., Xiang, H., Severinov, K., Zhang, X. and Wang, Y. (2019) Structure Studies of the CRISPR-Csm Complex Reveal Mechanism of Co-transcriptional Interference. Cell, 176, 239–253 e216.

20. Guo, M., Zhang, K., Zhu, Y., Pintilie, G.D., Guan, X., Li, S., Schmid, M.F., Ma, Z., Chiu, W. and Huang, Z. (2019) Coupling of ssRNA cleavage with DNase activity in type III-A CRISPR-Csm revealed by cryo-EM and biochemistry. Cell Res, 29, 305–312.

21. Smith, E.M., Ferrell, S., Tokars, V.L. and Mondragon, A. (2022) Structures of an active type III-A CRISPR effector complex. Structure, 30, 1109–1128 e1106.

22. Brouns, S.J., Jore, M.M., Lundgren, M., Westra, E.R., Slijkhuis, R.J., Snijders, A.P., Dickman, M.J., Makarova, K.S., Koonin, E.V. and van der Oost, J. (2008) Small CRISPR RNAs guide antiviral defense in prokaryotes. Science, 321, 960–964.

23. Athukoralage, J.S. and White, M.F. (2022) Cyclic Nucleotide Signaling in Phage Defense and Counter-Defense. Annu Rev Virol, 9, 451–468.

24. Steens, J.A., Zhu, Y., Taylor, D.W., Bravo, J.P.K., Prinsen, S.H.P., Schoen, C.D., Keijser, B.J.F., Ossendrijver, M., Hofstra, L.M., Brouns, S.J.J. et al. (2021) SCOPE enables type III CRISPR-Cas diagnostics using flexible targeting and stringent CARF ribonuclease activation. Nature communications, 12, 5033.

25. Santiago-Frangos, A., Hall, L.N., Nemudraia, A., Nemudryi, A., Krishna, P., Wiegand, T., Wilkinson, R.A., Snyder, D.T., Hedges, J.F., Cicha, C. et al. (2021) Intrinsic signal amplification by type III CRISPR-Cas systems provides a sequence-specific SARS-CoV-2 diagnostic. Cell Rep Med, 2, 100319.

26. Wang, L., Mo, C.Y., Wasserman, M.R., Rostol, J.T., Marraffini, L.A. and Liu, S. (2019) Dynamics of Cas10 Govern Discrimination between Self and Nonself in Type III CRISPR-Cas Immunity. Mol. Cell, 73, 278–290 e274.

27. Bokori-Brown, M., Martin, T.G., Naylor, C.E., Basak, A.K., Titball, R.W. and Savva, C.G. (2016) Cryo-EM structure of lysenin pore elucidates membrane insertion by an aerolysin family protein. Nature communications, 7, 11293.

28. Zivanov, J., Nakane, T., Forsberg, B.O., Kimanius, D., Hagen, W.J., Lindahl, E. and Scheres, S.H. (2018) New tools for automated high-resolution cryo-EM structure determination in RELION-3. eLife, 7.

29. Jumper, J., Evans, R., Pritzel, A., Green, T., Figurnov, M., Ronneberger, O., Tunyasuvunakool, K., Bates, R., Zidek, A., Potapenko, A. et al. (2021) Highly accurate protein structure prediction with AlphaFold. Nature, 596, 583–589.

30. Mirdita, M., Schutze, K., Moriwaki, Y., Heo, L., Ovchinnikov, S. and Steinegger, M. (2022) ColabFold: making protein folding accessible to all. Nat Methods, 19, 679–682.

31. Pettersen, E.F., Goddard, T.D., Huang, C.C., Meng, E.C., Couch, G.S., Croll, T.I., Morris, J.H. and Ferrin, T.E. (2021) UCSF ChimeraX: Structure visualization for researchers, educators, and developers. Protein Sci, 30, 70–82.

32. Liebschner, D., Afonine, P.V., Baker, M.L., Bunkoczi, G., Chen, V.B., Croll, T.I., Hintze, B., Hung, L.W., Jain, S., McCoy, A.J. et al. (2019) Macromolecular structure determination using X-rays, neutrons and electrons: recent developments in Phenix. Acta Crystallogr D Struct Biol, 75, 861–877.

33. Pantelic, R.S., Meyer, J.C., Kaiser, U., Baumeister, W. and Plitzko, J.M. (2010) Graphene oxide: a substrate for optimizing preparations of frozen-hydrated samples. J Struct Biol, 170, 152–156.

34. Danev, R., Buijsse, B., Khoshouei, M., Plitzko, J.M. and Baumeister, W. (2014) Volta potential phase plate for in-focus phase contrast transmission electron microscopy. Proc Natl Acad Sci U S A, 111, 15635–15640.

35. Zhang, J., Rouillon, C., Kerou, M., Reeks, J., Brugger, K., Graham, S., Reimann, J., Cannone, G., Liu, H., Albers, S.V. et al. (2012) Structure and mechanism of the CMR complex for CRISPR-mediated antiviral immunity. Molecular Cell, 45, 303–313.

36. Sofos, N., Feng, M., Stella, S., Pape, T., Fuglsang, A., Lin, J., Huang, Q., Li, Y., She, Q. and Montoya, G. (2020) Structures of the Cmr-beta Complex Reveal the Regulation of the Immunity Mechanism of Type III-B CRISPR-Cas. Mol. Cell, 79, 741–757 e747.

37. Zhang, J., Rouillon, C., Kerou, M., Reeks, J., Brugger, K., Graham, S., Reimann, J., Cannone, G., Liu, H., Albers, S.V. et al. (2012) Structure and mechanism of the CMR complex for CRISPR-mediated antiviral immunity. Mol Cell, 45, 303–313.

38. Mariani, V., Biasini, M., Barbato, A. and Schwede, T. (2013) lDDT: a local superposition-free score for comparing protein structures and models using distance difference tests. Bioinformatics, 29, 2722–2728.

39. Yamashita, K., Palmer, C.M., Burnley, T. and Murshudov, G.N. (2021) Cryo-EM single-particle structure refinement and map calculation using Servalcat. Acta Crystallogr D Struct Biol, 77, 1282–1291.

40. Casanal, A., Lohkamp, B. and Emsley, P. (2020) Current developments in Coot for macromolecular model building of Electron Cryo-microscopy and Crystallographic Data. Protein Sci, 29, 1069–1078.

41. Sokolowski, R.D., Graham, S. and White, M.F. (2014) Cas6 specificity and CRISPR RNA loading in a complex CRISPR-Cas system. Nucl. Acids Res., 42, 6532–6541.

42. Zhang, J., Graham, S., Tello, A., Liu, H. and White, M.F. (2016) Multiple nucleic acid cleavage modes in divergent type III CRISPR systems. Nucl. Acids Res., 44, 1789–1799.

43. Kazlauskiene, M., Tamulaitis, G., Kostiuk, G., Venclovas, C. and Siksnys, V. (2016) Spatiotemporal Control of Type III-A CRISPR-Cas Immunity: Coupling DNA Degradation with the Target RNA Recognition. Mol. Cell, 62, 295–306.

44. Johnson, K., Learn, B.A., Estrella, M.A. and Bailey, S. (2019) -Target sequence requirements of a type III-B CRISPR-Cas immune system. J. Biol. Chem., 294, 10290–10299.

45. Rouillon, C., Athukoralage, J.S., Graham, S., Gruschow, S. and White, M.F. (2018) Control of cyclic oligoadenylate synthesis in a type III CRISPR system. eLife, 7, e36734.

46. Hale, C.R., Zhao, P., Olson, S., Duff, M.O., Graveley, B.R., Wells, L., Terns, R.M. and Terns, M.P. (2009) RNA-guided RNA cleavage by a CRISPR RNA-Cas protein complex. Cell, 139, 945–956.

47. Tamulaitis, G., Kazlauskiene, M., Manakova, E., Venclovas, C., Nwokeoji, A.O., Dickman, M.J., Horvath, P. and Siksnys, V. (2014) Programmable RNA shredding by the type III-A CRISPR-Cas system of Streptococcus thermophilus. Mol. Cell, 56, 506–517.

48. Staals, R.H., Zhu, Y., Taylor, D.W., Kornfeld, J.E., Sharma, K., Barendregt, A., Koehorst, J.J., Vlot, M., Neupane, N., Varossieau, K. et al. (2014) RNA targeting by the type III-A CRISPR-Cas Csm complex of Thermus thermophilus. Mol. Cell, 56, 518–530.

49. Hale, C.R., Cocozaki, A., Li, H., Terns, R.M. and Terns, M.P. (2014) Target RNA capture and cleavage by the Cmr type III-B CRISPR-Cas effector complex. Genes & Development, 28, 2432–2443.

